# CoPPIs Algorithm: A Tool to Unravel Protein Cooperative Strategies in Pathophysiological Conditions

**DOI:** 10.1101/2024.12.11.627896

**Authors:** Andrea Lomagno, Ishak Yusuf, Gabriele Tosadori, Dario Bonanomi, Pierluigi Mauri, Dario Di Silvestre

**Author notes:** **Correspondence:** Dario Di Silvestre, Clinical Proteomics Laboratory, Elixir Infrastructure, Institute for Biomedical Technologies - National Research Council, F.lli Cervi 93, 20054 Segrate, Milan, Italy. **Funding information:** PRIN2022: 2022Z2TE5P and 2022LNHZAP; PRIN PNRR: P2022LY3F4.

## Abstract

We present here the Co-expressed Protein-Protein Interactions (CoPPIs) algorithm. In addition to minimizing correlation-causality imbalance and contextualizing PPIs to the investigated systems, it combines PPIs and protein co-expression networks to identify differentially correlated functional modules. To test CoPPIs, we processed a set of proteomic profiles from different brain areas of controls and subjects affected by idiopathic Parkinson’s disease or carrying a GBA1 mutation. Its robustness was supported by the extraction of functional modules, related to translation and mitochondria, whose involvement in PD pathogenesis is well documented. Furthermore, the selection of hubs and bottlenecks from the weighted PPI networks provided molecular clues consistent with the PD pathophysiology. Of note, like quantification, the CoPPIs algorithm revealed less variations when comparing disease groups than when comparing diseased and controls. However, correlation and quantification results showed low overlap, suggesting the complementarity of these measures. An observation that opens the way to a new investigation strategy that takes into account not only protein expression, but also the level of coordination among proteins that cooperate to perform a given function.

## 1 INTRODUCTION

The development of data-driven systems biology approaches is receiving a major boost from the improvement of-omics technologies, including single-cell ones, which allow for increasingly accurate large scale qualitative and quantitative measurements [1]. At the same time, the identification and validation of protein-protein interactions (PPIs) is benefiting from the development of new strategies and methodologies [2, 3, 4, 5]. Following these advances, network models are more and more being adopted to analyze complex biological systems by investigating the molecular relationships that characterize the network structure [6, 7, 8]; thus, the emergent properties that arise from it [9].

In the context of molecular network analysis, co-expression models have historically been used to describe large scale gene expression data through correlation patterns [10]. In particular, to identify modules of co-expressed genes related to phenotypic traits, pathophysiological states, treatments etc. [11]. For this purpose, the use of large-scale proteomic data may still be considered in its infancy. However, a growing number of studies are modeling the proteome as co-expression network [7, 12, 13, 14, 15, 16, 17, 18, 19, 20, 21, 22]. Indeed, the identification and quantification of the entire proteome is now possible [23]. In addition, the increasingly large number of samples, deposited in dedicated databases can be analyzed sistematically [24]. These two key factors offering an unprecedented opportunity for extensive evaluation of proteomic data through correlational studies [6].

So far, most studies combining large-scale proteomic data and co-expression network models have relied on weighted gene co-expression network analysis (WGCNA) [25]. The growing interest in this field has fueled the implementation of computational tools, including JUMPn [26] and ProtExA [27], to support differential expression, functional analysis, and identification of protein-protein interactions (PPIs) and co-expression network modules. Additionally, more recently, Buljan et al. developed an algorithm that, through the systematic measurement of the ratio between protein complex subunits, aims to identify compromised interactions [28]. However, prior to Buljan et al., we combined co-expression network models and PPIs to identify protein complexes, pathways and biological processes that showed altered correlation values in adipose tissue from patients with amyloidosis [7].

To automate the extraction of differentially correlated functional modules in pairwise comparisons, i.e. healthy vs. control, treated vs. untreated, etc., we propose here an algorithm called CoPPIs (co-expressed protein-protein interactions). Unlike other methods, we focus a priori on proteins that are known to interact physically or functionally to carry out a biological process (**Figure 1**). By focusing on PPIs, we aim to minimize the imbalance between correlation determinations and causality assessments [29]. On the other hand, this strategy is based on the assumption that, in order to cooperate, two or more proteins must be close and coordinated. In a way, this brings us back to starlings (*Sturnus vulgaris*) when they draw sinuous shapes in the sky as a result of a collective behaviour [30]. In other words, we assume that the execution of a given process or function requires that the cooperating subunits/proteins be expressed in a well-defined stoichiometry [7, 28, 31].

**FIGURE 1.**
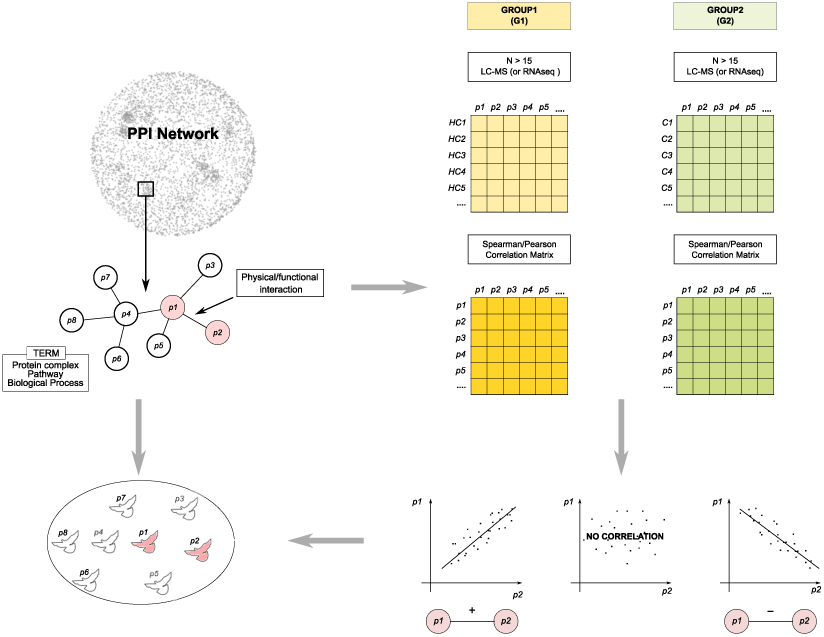
Genesis and objectives of the combination of PPI and coexpression models. This strategy aims to assess whether proteins interacting physically and/or functionally, in order to cooperate to carry out functions and processes, are coordinated. In some ways we took inspiration from starlings (*Sturnus vulgaris*) when they defend themselves from aerial predators [30]. In particular, a single starling observes its neighbor and imitates it. Moreover, it keeps a fixed number of its neighbors under control, about 7-8. Therefore, we assume that proteins can have a similar behaviour.

To test CoPPIs, and the robustness of the score it produces, we processed a set of proteomic profiles previously characterized by a collection of brain tissues from healthy controls and subjects affected by idiopathic Parkinson’s disease (IPD) or carrying a GBA1 mutation (PD-GBA1) [32]. Specifically, data from four brain areas, such as Substantia Nigra (SN), Striatum (STR), Occipital Cortex (OCC) and Middle Temporal Gyrus (MTG), were considered. In addition to provide a set of functional modules whose level of correlation significantly varies between conditions or brain regions, the correlation between protein pairs was used in the reconstruction of weighted PPI network models for a more accurate extraction of hubs and bottlenecks [6]. Unweighted PPI network models are in fact usually made of cell- and condition-type independent PPIs, thus their weighting is a way to contextualize them to the actual system investigated [33].

## 2 RESULTS

### 2.1 Co-expressed Protein-Protein Interactions (CoPPIs) algorithm flowchart

CoPPIs algorithm aims to automate the extraction of significant differentially correlated functional modules i.e. protein complexes, pathways, biological processes etc., by combining PPI networks with co-expression models inferred by LC-MS experimental data (**Figure 2A**). Starting from a protein data matrix *m*x*n* (where *m* represents genes/proteins and *n* samples) reporting semi-quantitative values (PSMs, Peak Area, Intensity), the first step involves the computation of *Spearman*’s correlation for pairs of identified proteins; alternatively, if data are normally distributed, *Pearson*’s correlation is applied. The same set of proteins (identification frequency (IF)=100% in all conditions) is processed for each investigated group of samples (e.g. healthy, diseased, treated).

**FIGURE 2.**
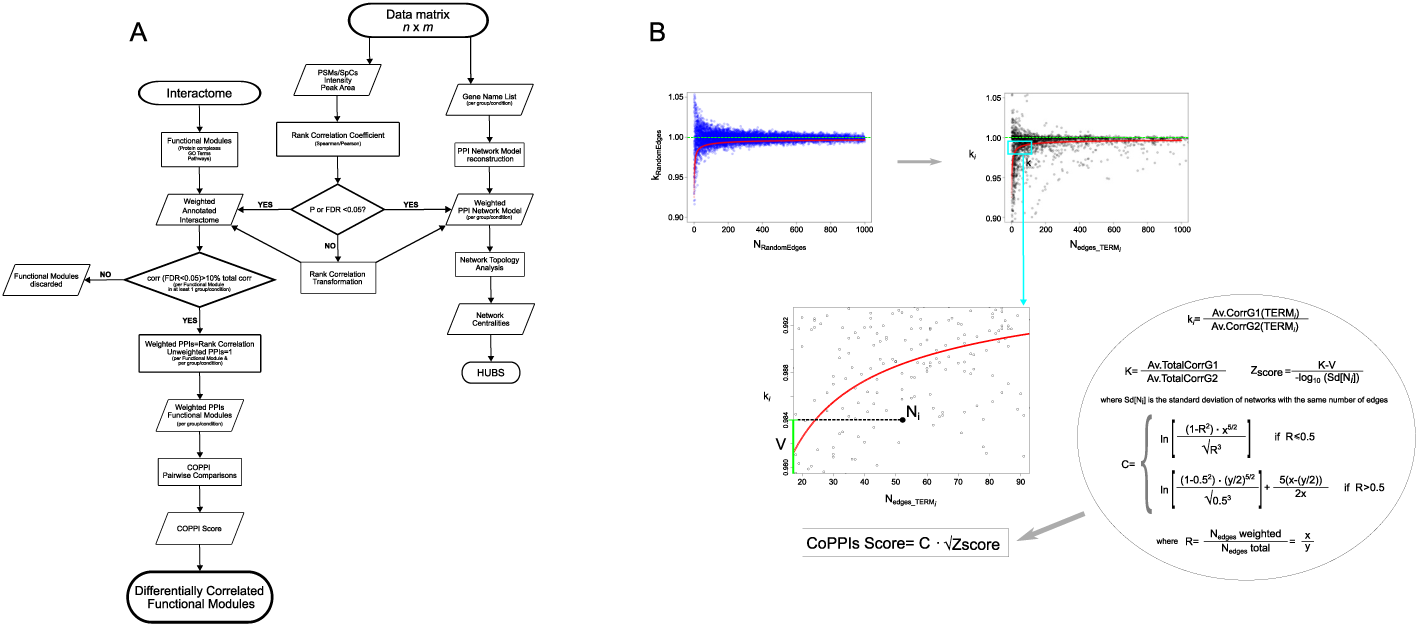
CoPPis algorithm. A) Flowchart showing the main steps of CoPPIs algorithm. B) Maths rationale of CoPPIs score. Top left, distribution of k_RandomEdges_ ratios between the average values from a number of randomly selected correlations (N_RandomEdges_) in a pair of groups/conditions compared; the distribution of k_RandomEdges_ ratios, which tends to K (green dashed line), is depicted as a function of the number of random correlations selected. Top-right, distribution of ratios between the average values of correlations k*_i_* corresponding to a specific biological term (N_edges_TERM_*_i_*) in a pair of groups/conditions compared; the distribution of k*_i_* ratios, which tends to K (green dashed line), is depicted as a function of the number of interactions of each term. In the box below, it is shown a detailed view of a generic term (N_edges_TERM_*_i_*) with ratio V, while inside the circle the mathematical steps that lead to the formulation of the CoPPIs algorithm score.

CoPPIs relies on PPI network models retrieved from STRING database. The acquired data is then transformed by a CoPPIs function into a graph, where proteins represent nodes, and protein-protein interactions form edges. The reconstructed interactome is matched with the *Spearman*’s correlation score measured per pair of identified proteins, and the corresponding edges are thus weighted. In this way, the condition-specific correlations are encapsulated in the PPI network model, providing a more refined representation of biological relationships. Additionally, non-significant correlations (FDR*≥* 0.05) undergo a process of transformation to be further penalized in comparison to their significant counterparts (**Supplementary Information**).

Once the interactome has been weighted with the transformed correlation scores, it is processed at a functional level by CoPPIs. This step allows the annotation of functional modules by referring to different sources, including GO Terms, Reactome, KEGG, Wikipathways, and CORUM [34]. Each module is represented as a graph, where the annotated proteins serve as nodes, and the edges correspond to PPIs weighted as previously described. Nodes that lack any connections, due to the absence of known interactions, are subsequently removed from the graph. For each module, unweighted edges, due to the lack of matching proteins in the input data matrix, are weighted to 1 by default. This step helps to reduce the extraction of false positive differentially correlated modules, mainly in the case of large ones where a low percentage of weighted edges could bias the detection of significant differences. Finally, the functional modules are processed in pairwise comparisons (e.g. healthy vs disease, treated vs untreated, WT vs mutant, etc.) and for each of them the CoPPIs score (**Supplementary Information**) is calculated to extract the significantly differentially correlated ones. (**Figure 2B**). The CoPPIs score is defined by a formula that increases with the ratio R, which is calculated as the number of significant edges for a biological term (x) divided by the total number of PPIs for the same biological term (y). To maintain an equal behavior for R lesser or greater than 0.5, two formulas were used, with equal partial derivatives with respect to x and y. The formula are designed so that, for a given value of R, a higher score is assigned when both x and y are larger and the partial derivatives with respect to x and y are both positive.

In parallel, a weighted PPI network model per group/condition is built and topologically evaluated by network centralities for the selection of hub and bottleneck nodes.

### 2.2 Protein correlation and proximity effect

Before calculating the CoPPI score comparing control subjects with those affected by IPD and PD-GBA1 across different brain areas, we investigated the hypothesis that protein correlation is influenced by a proximity effect, both at the level of physically interacting proteins and of subcellular localization.

Globally, we found that, compared to non-PPIs, PPIs were associated with higher mean correlation values (**Figure 3A**). This was further confirmed for all brain areas, where we also assessed a higher number of significant correlations associated with PPIs (**Figure S1**). Of note, the largest difference in correlation was observed for SN (**Figure S1A**), the main brain area involved in Parkinson’s disease [35].

**FIGURE 3.**
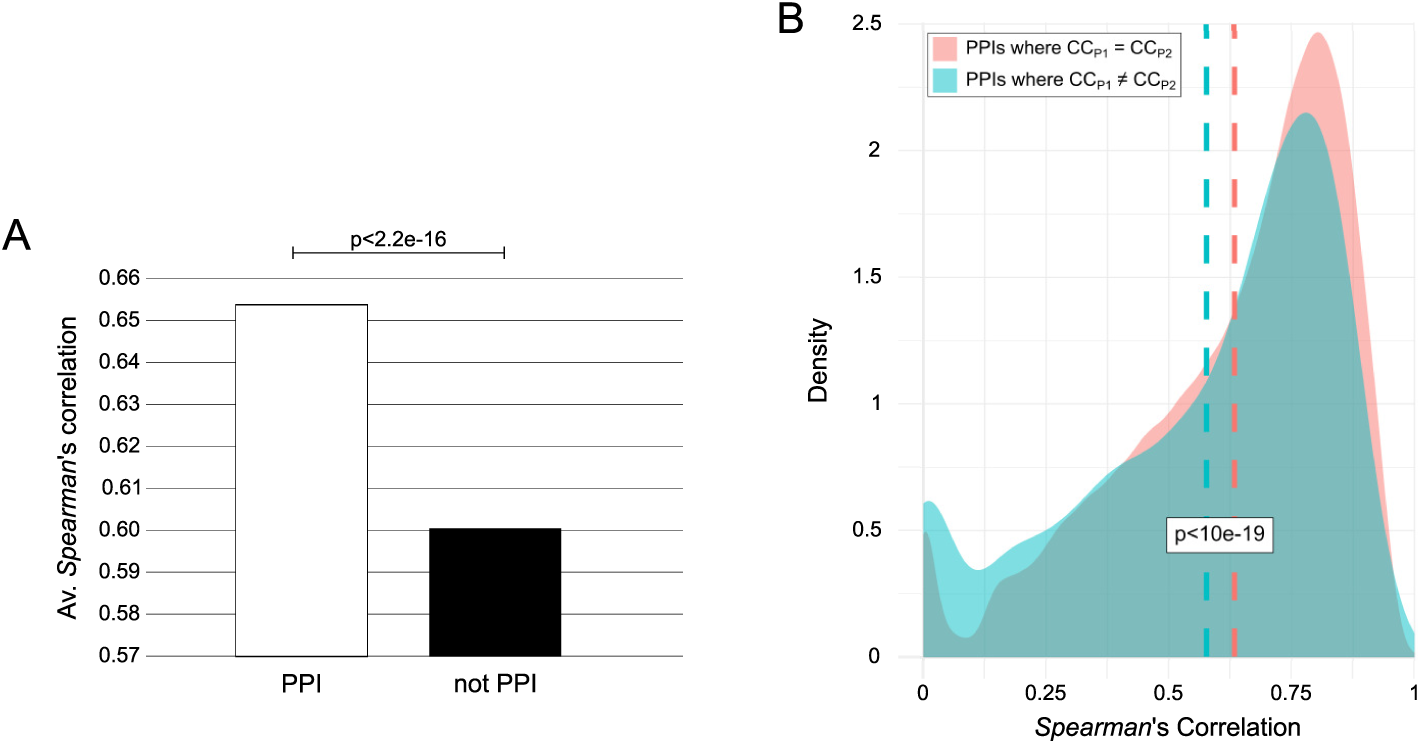
PPIs vs not PPIs correlation. A) Barplot reporting the average absolute values of significant correlations associated with protein pairs physically/functionally interacting (PPIs) and not (non-PPIs). B) *Spearman*’s correlation density distribution by comparing PPIs where P_1_ and P_2_ are annotated with the same Cellular Component (CC), and PPIs where P_1_ and P_2_ are annotated with a different Cellular Component (CC); blue and pink dotted lines indicate the average correlation. For both graphs, p value from *Student*’s *t*-test with NULL hypothesis that the mean of the distributions are the same.

Considering only PPIs, we evaluated the percentage of GO terms and pathways whose corresponding proteins were significantly correlated. Following these criteria, Compartments and Cellular Components (CC) had the highest percentage of terms with a significant correlation (**Figure S1**). In contrast, Wikipathways showed the lowest one. Taking Cellular Components as a reference, we found that PPIs consisting of protein pairs with the same subcellular localization showed a higher average correlation value than PPIs consisting of proteins localized in different compartments (**Figure 3B**). These results suggest a proximity effect on protein correlation, as well as further confirming a greater correlation between physically interacting proteins. Indeed, both Compartments and Cellular Components include protein complexes, whose correct functioning requires a well-defined stoichiometry between the subunits that compose them [36].

### 2.3 Differentially correlated functional modules in brain regions of control, IPD and PDGBA1 subjects

To test CoPPIs, we processed a collection of protein profiles obtained by analyzing different brain areas of Control, IPD and PD-GBA1 subjects [32]. For each group, we considered from 19 to 21 subjects, and for each subject we took into consideration the Substantia Nigra (SN), the Striatum (STR), the Occipital Cortex (OCC) and the Middle Temporal Gyrus (MTG). In particular, CoPPIs was used for evaluating the shift of correlation, by comparing the same brain area among the groups of subjects considered (**Table S1**, **Figure S2-S4**). Also, by comparing different brain areas within the same group (**Table S2**, **Figure S5-S10**).

Globally, most differentially correlated functional modules in intergroup comparisons were involved in mitochon-drial ATP synthesis, including the respiratory chain complexes, and neurotransmitter release, including clathrin-sculpted GABA and glutamate transport vesicle (**Figure 4**). Mitochondrial dysfunctions are strongly implicated in the etiology of idiopathic and genetic PD [37]; in particular, through the role of oxidative phosphorilation, complex I and Akt signaling pathway [38]. On the other hand, alterations in GABAergic and glutamatergic neurotransmission were associated with some PD symptoms [39].

**FIGURE 4.**
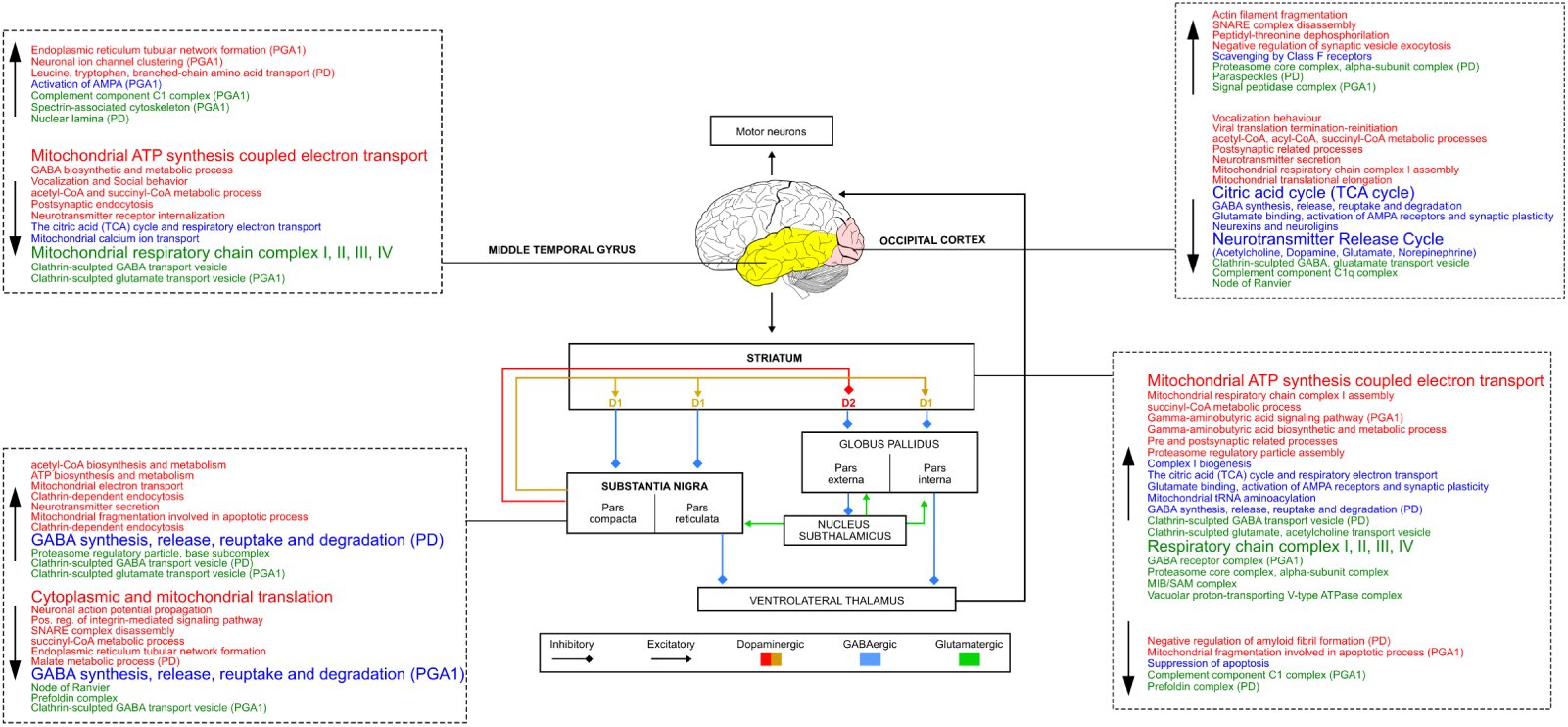
Scheme summarizing the Nigrostriatal pathway and the functional modules differentially correlated in the corresponding brain areas (SN, STR, OCC, MTG). Dashed boxes show the main Cellular Components (green), Biological Processes (red) and Reactome pathways (blue) whose correlation significantly varies by comparing Control *vs* IPD and PD-GBA1 subjects. In each box, up- and down-arrow indicates modules whose correlation is increased and decreased in IPD and PD-GBA1 (vs Control), respectively.

Similarly to label-free quantification (LFQ), the number of differentially correlated functional modules decreased when comparing the two groups of disease subjects, such as IPD and PD-GBA1 (**Figure 5A-B**). Nevertheless, quantitative analysis and correlation resulted complementary. Indeed, the results obtained following these approaches were barely overlapped. In that cases, we didn’t observe a relationship between the expression and correlation trend; some of them agreed, while others didn’t. SN and STR were the brain areas with the higher number of functional modules with an opposite trend. Of note, most of them were up-regulated in C and most correlated in IPD and PD-GBA1 (**Figure 5C**).

**FIGURE 5.**
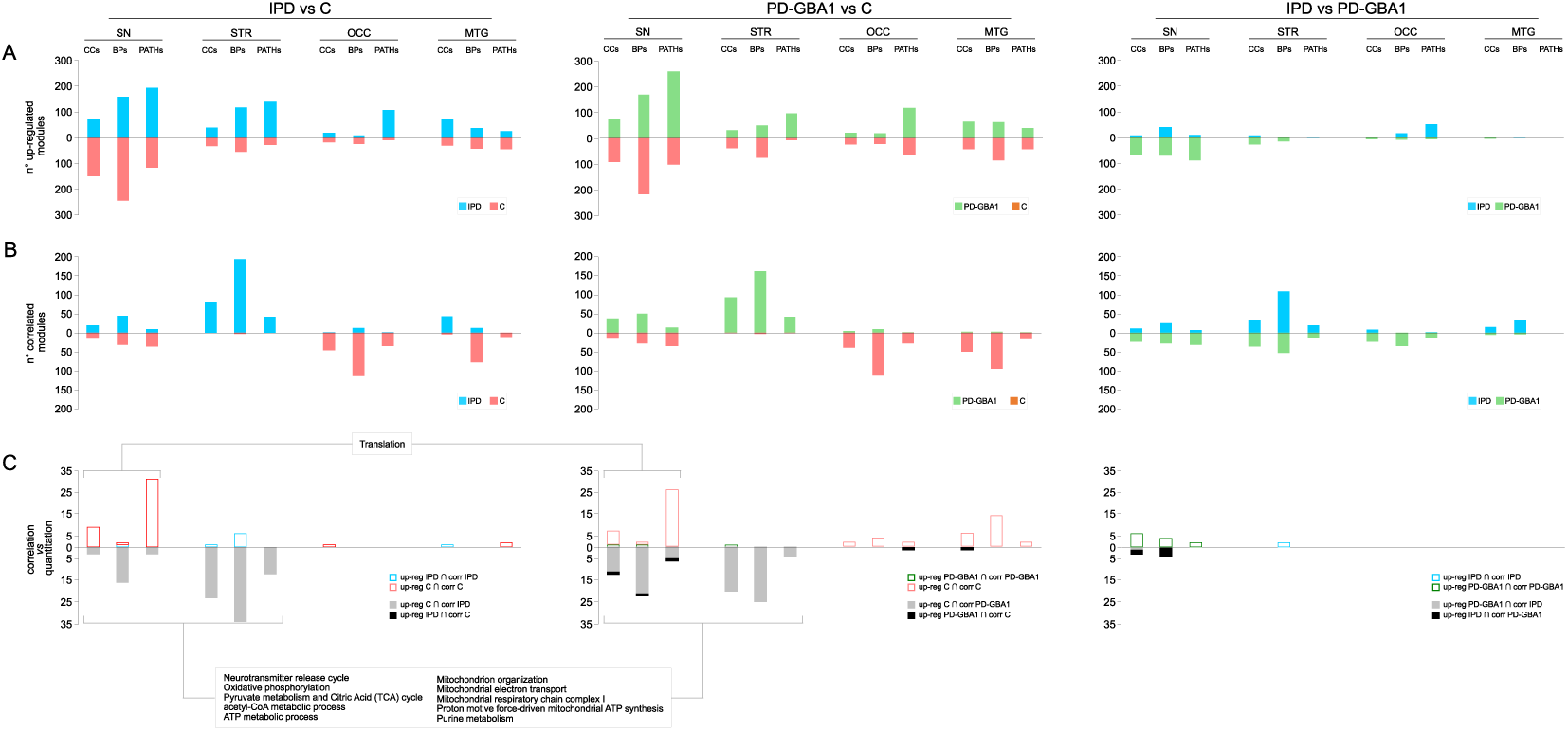
Histograms showing the number of A) differentially regulated and B) differentially correlated functional modules by comparing the same brain area in the following pairwise comparisons: IPD *vs* C, PD-GBA1 *vs* C and IPD *vs* PD-GBA1. In C), it is reported the number of functional modules with a similar (red, blue, green) and opposite (grey and black) trend of expression and correlation.

Due to the multitude of CoPPIs output, and the complexity of the study design, an overview of the main results by brain area is presented in the following paragraphs.

#### Substantia nigra

In SN of both IPD and PD-GBA1 subjects, CoPPIs detected a significant decrease of correlation in modules which refer to cytoplasmic and mitochondrial translation (**Figure 4**); noteworthy, through higher CoPPIs score values **Table S1**, **Figure S2-S4**). It represents a case where correlation and quantitation followed the same trend (**Table S3-S4 and Figure 5C**, **Figure 6**). Consistent with these results, down-regulation of both mitochondrial and cytoplasmic ribosomal subunits has been reported in the SN of PD patients, pointing to translation machinery impairment in disease pathogenesis [40]. Moreover, with the exception of IPD group, translation machinery-related modules were more correlated in SN vs all other brain areas, suggesting a specific role in this region (**Figure S8-S10**). This hypothesis is supported by a recent study showing an enrichment of translation-associated proteins in neuromelanin granules in the SN. These structures are formed after the inhibition of translation further indicating that this process is impaired in the SN of PD cases [41].

**FIGURE 6.**
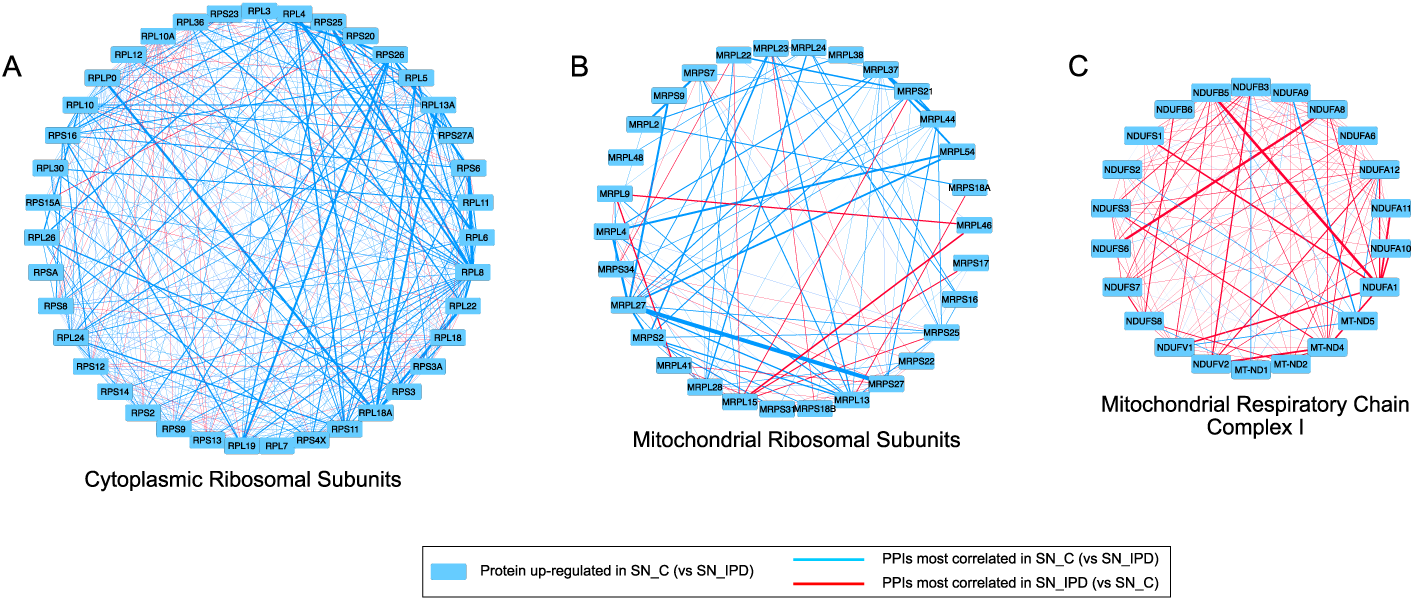
Functional modules showing the same and the opposite trend between quantitation and CoPPIs correlation. A) Cytoplasmic ribosomal subunits, B) Mitochondrial ribosomal subunits, and C) Mitochondrial respiratory chain Complex I in SN_C_ *vs* SN_IPD_; the thickness of blue and red edges is proportional to the correlation value.

Similar to the translation machinery, some functional modules related to ATP synthesis, mitochondrial electron transport and acetyl-CoA metabolism were down-regulated in patient groups, but with an opposite trend of correlation (**Figure 5C**). These quantitative variations observed following tissue analysis could mirror the mitochondrial dysfunctions occurring in PD, including a reduced bioenergetic capacity and the mitochondrial complex I deficiency [42]. They could also derive from the loss of functioning mitochondria and dopaminergic neurons, as a consequence of mitochondrial oxidative stress-mediated apoptosis [43, 44]. In agreement, in both patient groups, CoPPIs identified an higher correlation in mitochondrial fragmentation associated with apoptosis. This result aligns with the notion that, to mantain neural transmission in PD, the spared SN neurons enact a compensatory increase in mitochondrial density within the remaining axons and synapses [45]. Although this quantitative compensatory effect is not appreciated by classical quantitative proteomic approaches, it may be associated with the increased CoPPIs correlation in mitochondrial-related processes, including mitochondrial respiratory chain complex I (**Figure 6**), suggesting a protein coordination as part of such compensation.

Concerning other CoPPIs results from SN, the prefoldin complex resulted less correlated in both PD groups (**Figure 4**). Prefoldin is a molecular chaperone that co-localizes with *α*-synuclein in the lysosome. The knockdown of some Prefoldin subunits induced the accumulation of *α*-synuclein aggregates, suggesting a role of Prefoldin in preventing *α*-synuclein cell toxicity [46]. Finally, although at quantitative level IPD and PD-GBA1 group together, and their comparison provides a lower number of differentially expressed proteins and processes (**Figure 5A, Figure S11**), CoPPIs highlighted some differences supporting the heterogenous spectrum of parkinsonian disorders [47]. For instance, it is known that dopaminergic neuronal death in Substantia Nigra pars compacta (SNc) of PD patients affects GABAergic transmission in basal ganglia and this, in turn, augments GABAergic transmission in the Substantia Nigra reticulata (SNr) [48]. These pathological changes are consistent with our observation of increased correlation of GABA synthesis, vesicular transport release, reuptake and degradation in IPD. However, these functional modules were less correlated in PD-PGBA1 *vs* C (**Figure 4**), warranting further investigation.

#### Striatum

The increased GABAergic transmission from the basal ganglia to the SNr [48] may fit the increased correlation between proteins involved in functional modules linked to GABA synthesis, release, and metabolism. Striatal dopamine deficiency enhances activity in the ‘indirect pathway’ at the expenses of the ‘direct pathway’. This modulation is thought to increase GABA level in the striatum [49, 50]. Striatal dopamine denervation induces significant remodeling of corticostriatal and thalamostriatal glutamatergic synapses, which is consistent with increased synaptic transmission [51]. This hypothesis is supported by the increased correlation of glutamate binding, AMPA receptor activation and the synaptic plasticity pathway [52] and the clathrin-sculpted glutamate transport vesicle complex (**Figure 4**). The hypothesized increase in synaptic transmission could also explain the increased correlation of functional modules involved in ATP synthesis, essential to support the organization of synaptic vesicle pools, and neurotransmitter release during intense neuronal activity [53]. However, mitochondrial related processes were down-regulated. Finally, as demonstrated by Mitchell et al., endogenous glutamate can induce apoptosis of striatal projection neurons [54], a mechanism consistent with the decreased correlation between proteins involved in suppression of apoptosis.

#### Occipital cortex and Middle temporal gyrus

In contrast to SN and STR, in both OCC and MTG we recorded a reduced correlation between proteins involved in mitochondrial energy production, as well as neurotransmission, including GABA (**Figure 4**). In the OCC lobe of cognitively normal PD patients, Zhu et al. observed a small but statistically significant decrease in NAD and ATP concentration, suggesting that defects in energy metabolism are quantifiable even before regional neurological deficits become evident [55]. Meanwhile, reduced GABA levels were associated with visual hallucinations [56]. The reduced correlation between proteins involved in mitochondrial energy production, including ATP synthesis, TCA cycle, and respiratory channel complex I, II, II, and IV, was more consistent in MTG than in OCC. Regarding glucose metabolism, some authors have observed hypometabolism in several brain regions of PD patients [57], including the middle temporal gyrus [58], and associated this with the onset of significant cognitive decline. Although CoPPIs have not identified a differential correlation in modules related to glucose metabolism, glucose is a critical energy substrate for the brain, essential in ATP production, management of oxidative stress, and the synthesis of neurotransmitters, neuromodulators, and structural components [59].

### 2.4 Hubs and bottlenecks in weighted PPI networks

The correlation score calculated and transformed by CoPPIs was used to weight a PPI network model, per region and group, and extract topological relevant nodes, such as hubs and bottlenecks (**Table S5**). Both unweighted and weighted PPI network models showed scale-free distribution (**Figure S12**). From the comparison of the average centrality values, major differences were observed for Degree and Bridging (**Table1**); respectively, they were reduced and augmented in weighted models indicating a selection of the most relevant paths.

On average, brain areas from Controls showed a higher number of topologically relevant nodes (**Figure 7**). STR and MTG were the most enriched in hubs and bottlenecks in the control group, whereas SN showed these characteristics in IPD and PD-GBA1. Some proteins were hubs/bottlenecks in different brain areas, and OCC and MTG shared the highest number in both control and disease conditions. At the functional level, hubs/bottlenecks characterising conditions and brain areas were matched with the differentially correlated modules extracted by CoPPIs. For instance, the increased correlation of mitochondrial-related processes in SN and STR of patient groups matched with the selection of mitochondrial-related hubs/bottlenecks (**Table S6**). The same was observed in OCC and MTG areas of the control group (**Figure 4, Figure S13**).

**FIGURE 7.**
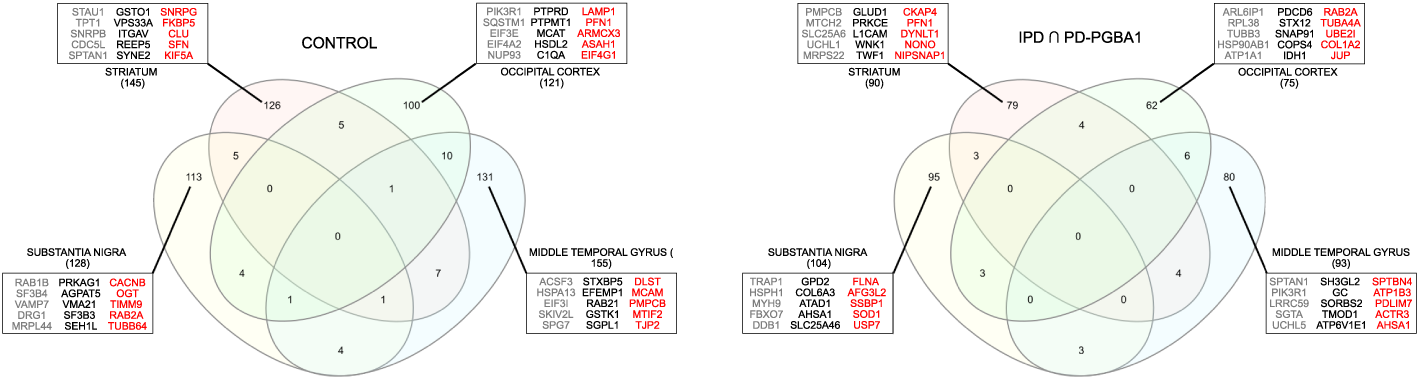
Venn diagram of hubs and bottlenecks found in brain areas from A) Controls and B) IPD and PD-GBA1 subjects. Boxes show the best-five ranked hubs (in grey, by betweenness and centroid), bottlenecks (in black, by betweenness and bridging) and hubs/bottlenecks (in red, by betweenness, centroid and bridging) per brain area (sorted by betweenness).

In the context of the highest-ranked topologically relevant nodes, proteins previously associated with Parkinson’s disease were identified (**Figure 7**). An overview by brain area is provided in the following paragraphs.

#### Substantia nigra

Among hubs/bottlenecks characterizing IPD and PD-GBA1 groups, we found Superoxide dismutase 1 (SOD1), an essential antioxidant enzyme, the heat shock protein 75 kDa (TRAP1) is a regulator of mitochondrial homeostasis, both implicated in protection against oxidative stress and mitochondrial dysfunction associated with PD pathogenesis [60, 61]. Noteworthy, two different SN hubs/bottlenecks, such as F-box only protein 7 (FBXO7) and Ubiquitin-specific-processing protease 7 (USP7), have been proposed to play a crucial role in the pathogenesis of PD. They are involved in regulating the balance between protein synthesis and degradation. Indeed, while FBXO7 is involved in the ubiquitination of various target proteins, UPS7 is a deubiquitinating enzyme. Moreover, UPS7 modulates the stability of FBXO7 and mitigate ER stress-induced cytotoxicity and apoptosis by preventing FBXO7 proteasomal degradation [62].

In the SN of control subjects, we found the O-Linked N-Acetylglucosamine (GlcNAc) Transferase (OGT) that catalyzes O-GlcNAcylation, a post-translational modification essential for the survival of dopaminergic neurons [63]. Another relevant hub characterizing this area/condition is the mitochondrial ribosomal protein L44 (MRPL44). In addition to further highlighting the role of the mitochondrial translation [40],this result is in line with evidence that the absence or defects in MRPL44 are associated with neurodegenerative diseases in humans, including PD [64].

#### Striatum

The best-ranked STR bottleneck of IPD and PD-GBA1 subjects resulted the glutamate dehydrogenase 1 (GLUD1), a mitochondrial matrix enzyme that catalyzes the oxidative deamination of glutamate to *α*-ketoglutarate which serves as a TCA cycle intermediate. GDH activity could support the mitochondrial factory for protecting neurons against metabolic failure [65]. Thus, its role could be consistent with an increased glutamate synaptic transmission (**Figure 4**) probably induced by striatal dopamine denervation [51]. Another bottlenecks, L1 protein (L1CAM), was already proposed as hub in human postmortem SN samples analyzed at transcriptomic level, and processed by weighted gene co-expression network analysis (WGCNA) [66]. Moreover, L1CAM is widely used as a target biomarker to detect Parkinson-associated neuronal EVs in human serum [67]. Ubiquitin carboxy-terminal hydrolase L1 (UCHL1) is instead a protein hub highly expressed in the brain where acts as a deubiquitinase enzyme. Although its association with PD as a genetic risk is controversial [68], UCHL1 Its interactors include proteins related to the development of PD, such as alpha-synuclein and ubiquitin-protein ligase parkin [69]. Finally, we found Nipsnap Homolog 1 (NIPSNAP1) in the double role of hub and bottleneck. Recent evidence demonstrated it is involved in regulation of mitophagy and probably can act as a sensor for mitochondrial health [70].

Glutathione s-transferase omega 1 (GSTO1) emerged as the most critical bottleneck in controls. In Drosophila it exerts a protective mechanism in mutant parkin restoring the activity and assembly of the mitochondrial F(1)F(0)-ATP synthase [71]. Finally Clusterin (CLU) has also been widely described as interfering with amyloid formation in many neurodegenerative disorders [72]. In particular, it binds -synuclein oligomers preventing an oligomer-induced increase in ROS production. Therefore, it suggests a neuroprotective role by suppressing -synuclein oligomer toxicity [73].

#### Occipital cortex

The Synaptosome Associated Protein 91 (SNAP1) was found as a bottleneck in IPD and PD-GBA1 subjects. Involved in the regulation of clathrin-dependent endocytosis, it has recently linked to PD [74]. Similarly, Junction Plakoglobin (JUP), which has the double role of hub and bottleneck has been associated with Parkinson’s disease initiation [75].

On the side of controls, we noted the presence of 3 different translation factors, such as Eukaryotic Translation Initiation Factor 3 Subunit E (EIF3E), Eukaryotic Translation Initiation Factor 4A2 (EIF4A2) and Eukaryotic Translation Initiation Factor 4 Gamma 1 (EIF4G1). Of note, genetic variant of EIF4G1 [76] and EIF3E [77] have been linked to PD, while EIF4A2 has been described in pediatric cases with developmental delay and dystonia-tremor syndrome [78]. Another interesting hub is Sequestosome-1 (SQSTM1) that, in combination with p62, promotes the degradation of unwanted molecules by macroautophagy, thereby serving as a signling hub for multiple pathways associated with neurodegeneration and making it a potential therapeutic target [79]. Deregulation of SQSTM1/p62 has been linked with a variety of neurodegenerative disorders, including PD. In the context of autophagosome/lysosome maturation and function, we found also acid ceramidase 1 (ASAH1)[80] and Lysosomal Associated Membrane Protein 1 (LAMP1). LAMP1 is a transmembrane glycoproteins localized in lysosomes and late endosomes. Several studies have confirmed that proteosomal and LAMP1-associated endolysosomal pathway dysfunctions lead to an increased toxicity of synuclein aggregates [81]. [82].

#### Middle temporal gyrus

Endophilin-A1 (SH3GL2) is the best ranked bottleneck found in IPD and PD-GBA1 subjects. It has been described among the genetic risk factors for Parkinson’s disease [83]. Specifically, SH3GL2 mutations are associated with the impairment in synaptic vesicle endocytosis prior to loss of dopaminergic neurons, therefore representing a significant contributor to PD pathogenesis [84].

As for controls, SPG7 Matrix AAA Peptidase Subunit, Paraplegin (SPG7) was selected as hub. SPG7 is associated with hereditary spastic paraplegia overlapping with mitochondrial disease features. In particular, SPG7 pathogenic variants impair mitochondrial homeostasis owing to mitochondrial DNA abnormalities, a possible link explaining the parkinsonism symptoms frequently observed in patients affected by this disease [85].

## 3 MATERIAL AND METHODS

### 3.1 CoPPIs algorithm

CoPPIs algorithm was built in R and different libraries, including *stringr, matrixStats, rbioapi, igraph, openxlsx, Hmisc, RCy3, tidyr* and *dplyr* were adopted. The complete code is available at https://github.com/lomi95/CoPPIs. CoPPIs uses PPI network models from STRING [86]. The current version of CoPPIs include the human interactome reconstructed by considering the interactions *experiments* (score *≥* 0.15) or *database* (score *≥* 0.3) annotated; of course, different interactomes and thresholds may be adopted.

### 3.2 Data test

To test the functionality of CoPPIs in identifying molecular alterations, we processed a set of proteomic profiles previously characterized by analyzing a collection of brain tissues from idiopathic Parkinson’s disease patients (IPD) and PD patients carrying a GBA1 mutation (PD-GBA1) [32]. The original collection counted 21 IPD patients, 21 PD-GBA1 patients and 21 controls from five different brain regions including the substantia nigra (SN), the striatum (STR), the occipital cortex (OCC), the middle temporal gyrus (MTG) and the cingulate gyrus (CG). However, due to a different number of available samples among control, IPD and PGBA1, the CG region was not considered (**Table S7**).

### 3.3 Label-free quantitation

The log2 intensities values of the identified proteins, already normalized to the total signal, were compared using a label-free quantification approach, as previously reported [7]. Specifically, the data matrix dimensionality (SN: 4534 proteins identified in 20 Controls, 20 PD and 20 PGBA1; STR: 4714 proteins identified in 19 Controls, 21 PD and 21 PGBA1; OCC: 4411 proteins identified in 21 Controls, 21 PD and 21 PGBA1; MTG: 5002 proteins identified in 21 Controls, 20 PD and 21 PGBA1) was reduced by linear discriminant analysis (LDA). Only proteins with P-value *≤*0.001 were retained. Pairwise comparisons were performed by DAve index [87]. All data processing were performed using a in house R script.

### 3.4 Topological analysis of weighted PPI network models

The network topological analysis was performed using an *in house* R script that calculate weighted betweenness, centroid, and bridging centralities on PPI network models, with correlation used as edge weight. Specifically, centroid formula refers to Gräßler et al. [88], while concerning bridging, instead of using the degree, the sum of incident edges per node was adopted [89]. The statistical significance of the model’s robustness was assessed by comparing the average weighted betweenness of our networks with the betweenness of random networks [90]; the random networks were generated by creating networks with the same degree distribution (using the *degseq(method = “vl”)* function from the *igraph* package) and assigning the original weights to the new edges, thereby preserving the identical weight distribution. Visualization was carried out using the *ggplot* package. Subsequently, nodes with both betweenness and centroid values above the 75th percentile were considered hubs, while nodes with both betweenness and bridging values above the 75th percentile were classified as bottlenecks.

## 4 DISCUSSION AND CONCLUSIONS

In this study we introduce CoPPIs alghoritm, a comprehensive end-to-end workflow that combines PPI models and experimental protein profiles to identify differentially correlated functional modules. The large-scale correlation assessment of PPI represents a novelty and fits a broad applicability to established and novel MS technologies. The results obtained in the testing phase suggest the general utility of the algorithm, demonstrating that it is complementary to classical protein quantification. Indeed, the results obtained from these two different approaches showed a low overlap. However, like quantification, CoPPIs extracted few significant differences when disease groups were compared. While a higher number emerged from the comparison between control and disease groups, highlighting the robustness of the CoPPI strategy. The quality of the results obtained by CoPPIs has been further supported by the extraction of functional modules, i.e. translation [91, 41, 40] and mitochondria-related [45, 43, 44, 42], whose involvement in Parkinson’s disease pathogenesis is widely documented. In this context, the increased correlation of mitochondrial processes, in the face of a decrease in quantity, suggests a stress response and coordination that could fit with a compensatory effect already proposed in previous works [45]. This brings us to the behaviour of starlings that draw their choreographed shapes in the sky in a stressful situation, that is, when they have to defend themselves from aerial predators [30]. These mechanisms are locally developed, and an individual bird observes its neighbours, about 7-8, and imitates them by following a single segment of the collective flight. Similarly, a proximity effect on protein correlation was observed, providing compelling evidence for the interdependence of protein interactions and their coordinated behaviour within distinct functional contexts. This is consistent with the correlation between protein localization and function; a phenomenon already emerged at the level of gene expression, where genes with similar expression patterns participate in the same signaling and regulatory pathways and circuits [92].

Since proteins represent the ultimate effector of biological functions, we expect that advanced PPI models, which clarify their relationships, will be powerful tools for deciphering the role of protein networks in physiology and disease. In addition to minimizing the imbalance between correlation determinations and causality assessments [29], the inclusion of protein correlation scores in PPIs aims to achieve a more accurate identification of protein hubs and bottlenecks. Indeed, up to today, PPI networks have been often matched with gene expression level, and weighted through correlation scores computed from gene expression profiles [6, 93]. On the other hand, besides signaling pathways, PPI network models are usually reconstructed from cell-type independent PPIs, which is in contrast to our understanding that protein dynamics are context-specific and highly dependent on their environment [33]. As a consequence, PPIs weighting with experimentally defined protein profiles represents a step forward for addressing these weaknesses and contextualize our models. This strategy produced network models devoid of potentially less relevant interactions, thereby more closely adhering to the biological systems under investigation. Here, we demonstrate that this fundamental improvement, combined with the identification of topologically relevant nodes, enables the discovery of molecular hits that align with the pathophysiology of PD [60, 61, 62, 63, 64, 66, 71, 73, 76, 77, 79, 81, 85].

In summary, the CoPPIs scoring system provides an elaborate assessment that considers both the magnitude of change and the structural characteristics of the interaction network. It is designed to avoid biases associated with pure ratio-based assessments and offers a robust metric to discern the functional relenance of differentially correlated terms. Conversely, it may suffer from a different sample size between the groups or conditions being compared. More generally, the sample size represents one of the major limitations to the effective application of the algorithm [94]. Indeed, correlational studies require a larger number of profiles than are tipically available in proteomics. However, with the rapid evolution of proteomic technologies, method standardization, and data archiving and cataloging, obtaining a sufficient number of proteomic profiles is becoming a realistic expectation in the near future. The data used in the testing phase in this study are moderately large; however the heterogeneity of groups and conditions facilitated a reliable evaluation of the algorithm’s behaviour. While these samples, collected post-mortem, allowed us to provide a snapshot of the late phase of the pathological process, the algorithm would be valuable to assess the co-expression variation at different disease stages. Furthermore, the analyzed protein profiles derive from a tissue analysis, making them a composite representation; for example, in the case of the SN, where different regions, such as the pars compacta (SNc) and reticulata (SNr), perform different functions [32]. Nevertheless, the results obtained by CoPPIs are very promising and improvements will be introduced in the near future, including capturing nonlinear relationships, the use of signaling networks and the integration of profiles that take into account post-translational modifications, such as phosphorylation [95]. These advancements hold great potential for further enhancing the accuracy of the CoPPIs algorithm and its ability to illuminate fundamental pathophysiological mechanisms that depend on complex protein networks.

**TABLE 1.**
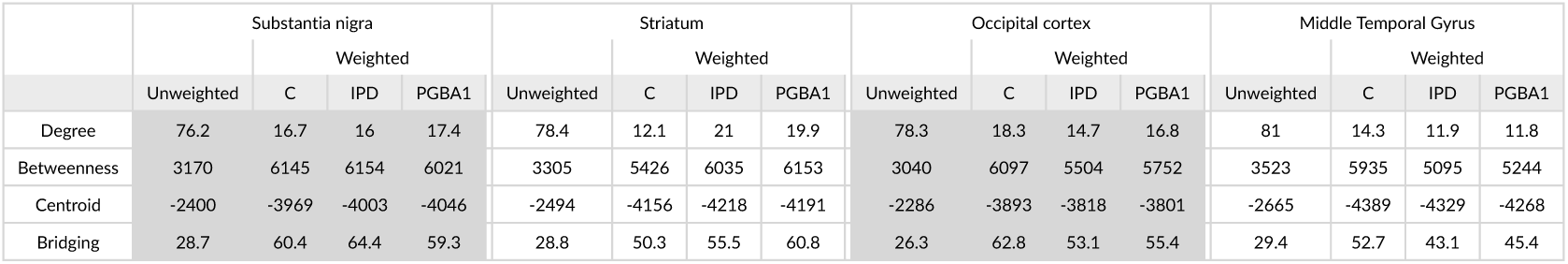
Weighted network topological analysis. Degree, Betweenness, Centroid and Bridging average values per brain area and condition are shown. Both unweighted and weighted PPI network models were analyzed.

## Abbreviations

CoPPIs: Co-expressed Protein-Protein Interactions
PD: Parkinson’s disease
WGCNA: weighted gene co-expression network analysis
PPIs: protein-protein interactions
IPD: idiopathic Parkinson’s disease
PD-GBA1: Parkinson’s disease with GBA1 mutation
C: control subjects
SN: Substantia Nigra
STR: Striatum
OCC: Occipital Cortex
MTG: Middle Temporal Gyrus
LC-MS: Liquid chromatography-Mass Spectrometry
PSMs: Peptide Spectrum Matches
IF: Identification Frequency
FDR: False Discovery Rate
GO: Gene Ontology
WT: Wild type
CC: Cellular Components
LFQ: label-free quantification
SNc: Substantia Nigra pars compacta
SNr: Substantia Nigra reticulata
EVs: extracellular vesicles
LDA: linear discriminant analysis.

## 5 ACKNOWLEDGEMENTS

We thank Valeria Bellettato for the technical and administrative support.

